# Within the fortress: A specialized parasite of ants is not evicted

**DOI:** 10.1101/002501

**Authors:** Emilia S. Gracia, Charissa de Bekker, Jim Russell, Kezia Manlove, Ephraim Hanks, David P. Hughes

## Abstract

Every level of biological organization from cells to societies require that composing units come together to form parts of a bigger unit (1). Our knowledge of how behavioral manipulating parasites change social interactions between social hosts is limited. Here we use an endoparasite to observe changes in social interactions between infected and healthy ants, using trophallaxis (liquid food exchange) and spatial data as proxies for food sharing and social segregation. We found no change in trophallaxis (p-value = 0.5156). By using K-function and nearest neighbor analyses we did see a significant difference in spatial segregation on day 3 (less than 8 millimeters; p-value < 0.05). These results suggest healthy individuals are unable to detect the parasite within the host.

## Introduction

Every level of biological organization from cells to societies require that composing units come together to form parts of a bigger unit (1). Where the composing units are themselves individuals, as in the case of social insects, the success of the group requires these units to be altruistic; explaining this behavior is conceptually challenging because one animal provides a benefit to another at a cost to itself. The seminal contribution of W.D. Hamilton (2, 3) provided a simple yet powerful framework for understanding altruism. This has since become known as Hamilton’s Rule and posits that a behavior or trait will be favored by selection, when *rb*-*c* > 0, where *c* is the fitness cost to the actor, *b* is the fitness benefit to the recipient, and *r* is their genetic relatedness. Since then numerous studies across a very diverse range of organisms from bacteria and yeasts to ants and mammals have demonstrated that the necessary conditions for altruism are upheld (4, 1).

Crucial to understanding the evolution of altruism has been determining how animals distinguish kin from non-kin because r must be >0 to satisfy Hamilton’s rule. An unlikely tool for studying altruism, it turns out, are parasites, the very antithesis of altruistic behavior. Parasites have evolved ways to break the code within kin groups, benefiting from their altruism, despite being completely unrelated to the donor (r = 0). The classical example is the Common Cuckoo (*Cuculus canorus*) that exploits the parental behavior of other bird species through egg mimicry and selfish behaviors by the chick. In societies such as ants, where altruism is expressed from sibling to sibling, diverse parasites ranging from other ants to beetles, flies, caterpillars and even mollusks have evolved ways to break the code to act like cuckoos (5,6). For example, caterpillars (i.e. *Maculinea rebeli*) perfectly mimic the chemical profile of larval ants and are carried into the nest by foraging workers, where they are then fed colony resources and consume the larval and egg stage ants (5,7,8,9). The general term for such organisms is social parasite and studying their chemical ecology and behavior has provided many insights into the mechanisms by which altruism works.

Although the color pattern of a cuckoo egg or the chemical cues of caterpillars entering ant nests are complex, it is conceptually easy to imagine how they evolved from ‘so simple a beginning’ (10). Indeed Darwin considered both cuckoo and socially parasitic ants, conjecturing that each emerged from non-parasitic progenitors (Chapter 7). More difficult to conceptualize is whether a parasite entering the body of an altruist can be recognized as a parasite since it is within the body of a colony member who presents kin recognition cues to the rest of the colony. Here test whether uninfected members of an ant colony can recognize siblings infected by a specialized endoparasite. We will use as our model the entomopathogenic fungus, *Ophiocordyceps unilateralis,* a highly specialized parasite of worker ants that manipulates host behavior to achieve transmission.

In recent years a number of studies have demonstrated that species within the complex *O. unilateralis s.l.* infect worker ants and adaptively manipulate host behavior by causing infected individuals to leave the colony and bite into vegetation before dying (11). The function of such manipulation is to provide a platform for spore release as post-mortem the fungus transitions from growing with the body to growing externally, forming a large stalk from which spores are produced and released onto the forest floor (12). Because the time from exposure and infection to behavioral manipulation is between 9 days and 3 weeks, during which time the infected ant is within the colony, this model offers the potential to examine how the colony responds to infected individuals. In this study we develop a system for within colony observation of infected and healthy individuals. We show that despite their status, infected ants are neither evicted from the colony nor prevented from leaving the nest to die and spread disease to their siblings.

## Results

### Infected ants receive food from siblings

We first set out to determine if infected ants received food from their siblings inside the nest area. We infected ten ants per colony in four different colonies of *Camponotus castaneus* species with a strain of *Ophiocordyceps unilateralis* fungus, which naturally infects this species in the wild (13). Ten ants were removed from the stock colony and injected with 1 µl *O. unilateralis* in solution with Grace’s insect media supplement with 10% fetal bovine serum (FBS). A further 10 ants were sham treated, and inject with 1 µl of Graces +FBS media. Both infected and sham treated ants were maintained together with 15 additional untreated individuals in a wooden chamber of volume 14.93 ± 0.53 cm^3^ placed within a cage of 451.61cm^2^ that served as a foraging area and contained sand. These ants were given water and 10% glucose *ad libitum.* We began continuous data recording from the third day post injection until day 18, moment at which there was the least amount of infected individuals alive. Ant behavior was recorded inside the nest with GoPro Heron 2 cameras for 24 hours/day. We then scored behavior from playback on screens. We first focused on food exchange between individuals since out hypothesis was that infected ants receive food at a different rate from non-infected ants. Worker ants cannot eat solid food but instead exchange liquids in a process called trophallaxis. We followed 17 focal individuals from one colony for a total observation 976.24 hours. Using a mixed effect linear model, programmed in R, we used trophallaxis duration as a function of day post infection and using ant identification as a random effect we found no significance *p-value* = 0.5156. We found no significant patterns in differences in either duration or count of trophallaxis (Figure 1). Our quantification of observations includes only within nest exchange of liquids. We did however also observe that ants infected by *O. unilateralis* would receive trophallaxis from nest mates when outside the nest. Individuals were even fed in the minutes and hours before they were behaviorally manipulated to ascend vegetation before biting bite into the twigs we provided and dying. We therefore found no evidence that infected ants were refused food from other colony members.

**Figure 1-.**
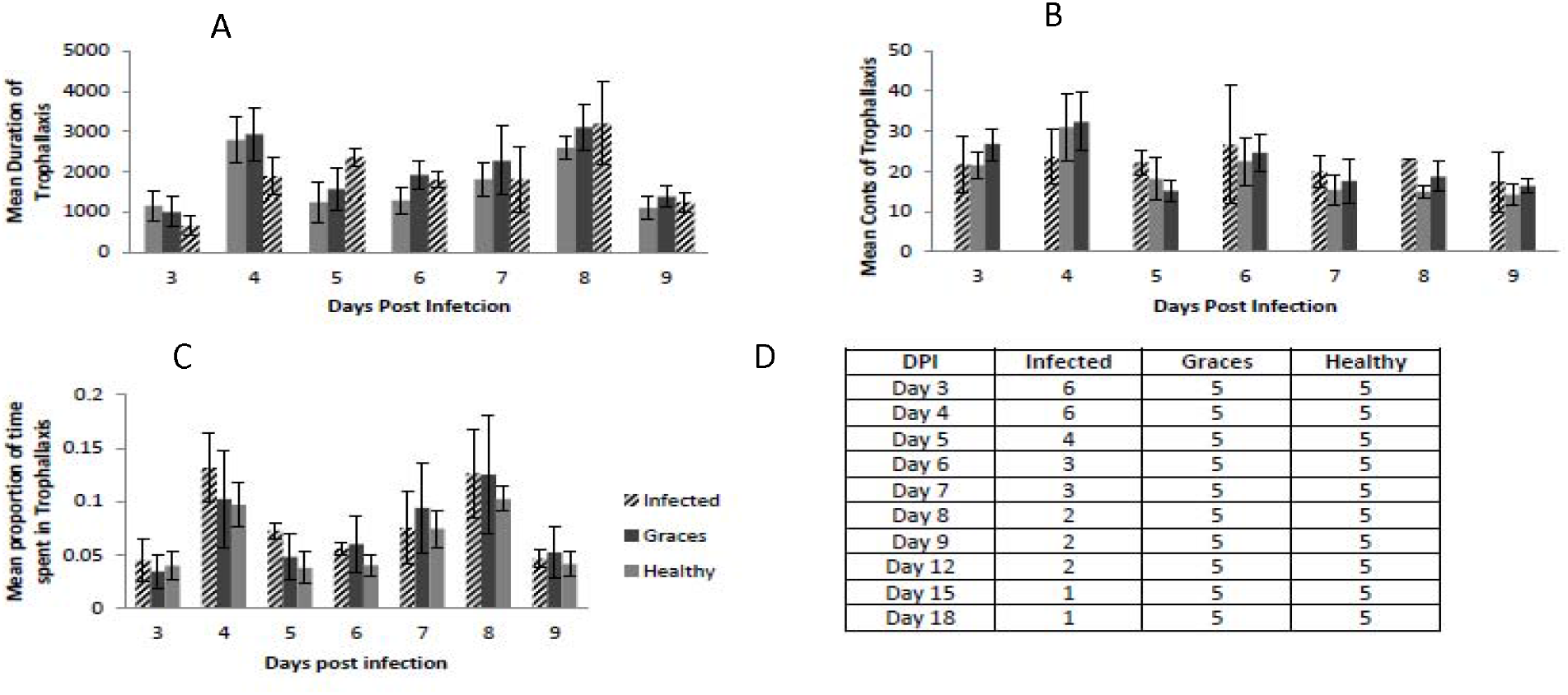
Trophallaxis data collected from videos on days 3-9, error bard represent the standard error of the data. We were unable to see any significance difference between treated individuals and healthy. (A) Shows no differences between infected, graces and healthy ants although we do see an interesting pattern of duration increase on day 4 and 8. (B) Mean count of trophallaxis changes slightly throughout the days we have observed. (C) The proportion of time spent in trophallaxis.

### Infected ants are not attacked by siblings

It might be expected that infected ants are attacked more often. Although an infected ant can only infect other ants following its own death and the subsequent growth of the stalk from its head, it is possible that the increase of fungal cells within the worker ants changes some aspect of it phenotype, such as smell, causing other ants to attack it. Because aggression might be rare and fleeting we observed the behavior of 2 colonies (30 ± 5 ants/colony) 24 hours/day for 18 days (still in progress). We saw no aggression between untreated control ants and either infected or sham treated ants. We conclude based on continuous observations over the entire course of infection that infected ants are not attacked by their siblings.

### Infected ants are initially distanced from colony members but this declines over time

Although we found no aggression there were subtle indications of infected ant segregation. To measure this we applied spatial point process approaches to within-nest ant locations. To examine spatial interaction behavior between healthy and infected ants, we examine the K-cross function (14) between healthy and infected ants, which is the expected number of infected ants that would be found within a distance *d* of each healthy ant.

Our null hypothesis is that ant infection status does not affect the tendency to group together or avoid other ants. We used a nonparametric permutation test (15) to test for significant deviation from this null behavior. In Figure 2 we show the observed K-cross function (in red) for all the data, as well as for days 3, 6, and 9, together with 1000 K-cross functions (in black) simulated from the null model by randomly permuting the labels (e.g., healthy, infected, or sham) of the ants 1000 times, and calculating the K-cross function between healthy and infected ants under each of these permutations. Significant deviation from the null model is indicated by an observed K-function that lies on the edges (tails) of the envelope of K-functions simulated under the null model, and empirical *p*-values can be computed by considering the rank of the observed K-function within the envelope. Significant deviation from null behavior only occurs day 3 at short spatial distances (less than 8 millimeters; p-value < 0.05), where the observed K-function lies on the lower tail of the permuted K-functions, suggesting spatial segregation at small spatial scales.

**Figure 2-.**
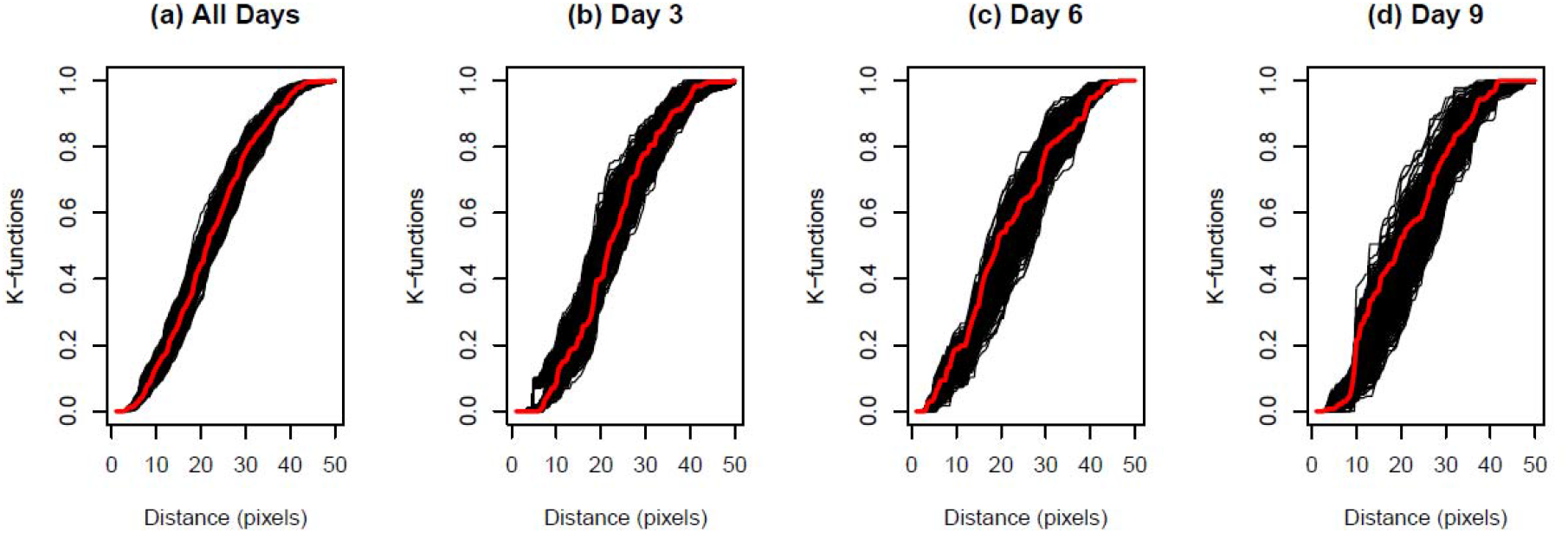
K function analysis.

To examine this potential small-scale ant interaction behavior between healthy and infected ants, we found the nearest neighbor to each ant at each time point. We then tested for deviation from the null assumption that ants are equally likely to have any other ant as nearest neighbor by again permuting the ant labels at each time point and recording the permuted label of each ant’s nearest neighbor. Table 1 shows the proportion of healthy ants with an infected nearest neighbor over all observed days, and for day 3, day 6, and day 9, together with the empirical *p*-value under the null, obtained using 1000 permutations of ant labels at each time point of observation. We see significant differences on day 3 and when we pool all the data together.

**Table 1-.** Nearest neighbor analysis we can see there is a significant difference between healthy and infected on when looking at all three days combined and only on day 3.

| Time | Proportion | Permutation Test p-value |
| --- | --- | --- |
| All Days | 0.110 | 0.002 |
| Day 3 | 0.108 | 0.004 |
| Day 6 | 0.208 | 0.48 |
| Day 9 | 0.104 | 0.24 |

## Discussion

Our data suggests no aggression towards infected individuals. We also see no distinct differences in the mean duration or counts of trophallaxis between infected and uninfected individuals (p-value = 0.5156). Our linear mixed effect model shows trophallaxis had no relation with treatment and trophallaxis is stochastic. Other studies have used trophallaxis as a tool to study social immunity, making similar observations to the ones we have made here, yet their results are an increase the amount of trophallaxis that occurs 24 hours after infection with a fungal pathogen (16, 17). An important factor these papers did not take into account is time, they only made observation 24 hours after the infection, this experiment on the other hand observed the ontogeny of behavior within the nest. By using a one chamber scenario in a cage where ants were able to freely move and interact with one another enables us to observe more naturalistic interactions that has been lacking in the ant-pathogen research.

Our data suggest there are no shifts in behavior towards infected individuals, suggesting healthy individuals are unable to detect *Ophiocordyceps* infection. The spatial point process analysis revealed that by and large there is no evidence for spatial segregation of infected ants. The only exception was the slight differences in spatial segregation between healthy individuals and those infected at small spatial scales on day 3 of the infection, but not on days 6 and 9. These minute changes in spatial arrangement could be caused by changes in individual infected ant behavior and are not likely to be indicators of social exclusion, which we would expect to increase in strength with time from infection. We did not test for any relationship between spatial segregation and the identity of the focal individuals in relation to who perform the most trophallaxis. Data collection for both trophallaxis and distance data were collected on Colony 2 the sample sizes we have may be masking the effect of *Ophiocordyceps* on the infected individuals.

Within nest distance observations has been done before by using images to determine spatial fidelity and time budgets of *Leptothorax acervorum* (18, 19), their observations did not take into account how pathogens may change social dynamics within a colony nor did they do continuous behavioral observations. We were able to observe rare interactions and behaviors that have previously not been described. Being able to follow individuals through time and space lends itself to be a powerful tool for further understanding the ontogeny of behavior within infected individuals. Although our trophallaxis and distance results were not significant we can still progress our understanding between uninfected individuals and those being parasitized.

In order to establish if these results are caused by the evolutionary history between *Ophiocordyceps* and host we should observe non-coevolved pathogen species. These behavioral assays enable us to further explore the role of parasites in not only the behavioral of the single host, but also in the colony host. The ability to combine behavioral observation and spatial dynamics as a tool to make very fine detailed observations enables us to further tease out the dynamics of the colony and those infected. Another powerful tool that we could add to this type of behavioral assay is chemical cues, such as cuticular hydrocarbons.

Chemical communication is the method of communication within an ant colony (20,21). Therefore individual odor changes could signify caste allocation (22) and colony members could also use it as a methods to determine health. Using continuous, detailed observations and cuticular hydrocarbons would give insight into how these infected individuals are perseived by their nest mates.

## Methods and Materials

### Ant collection and stock colony maintenance

Ants were collected in South Carolina during October 2012. Colonies were collected by following foragers to nest sites that were then dug up. Colony 1 consisted of sexuals, brood and about 120 workers; collected 10/4/2012. Colony 2 formed by 100 workers and brood; collected 10/5/2012. Colony 3 formed by100 workers and brood; collected 10/3/2012. Colony 4 has queen, brood and 120 workers. Colonies were maintained by providing them sugar water and water ad libitum and changed once a week. From these colonies we collected individuals to run our experiment on.

### Infection techniques

O. unilateralis Infections were done as described in de Bekker *et al*., 2014/submitted. Single fungal colonies were placed in a sterile 2 mL tube with two 8/32 inch metal balls (Wheels Manufacturing Inc.) and 200 µl Grace’s medium (Sigma) freshly supplemented with 10% FBS (PAA laboratories Inc.). The colony tissue was lysed using TissueLyser II (Qiagen) at room temperature for 60 sec. at 30 freq/sec. This processed enabled us to obtain single hyphae used at a mean concentration of 3.9 × 107+/−1.1 × 10^7^ hyphae/ml for infection. Infections were done by injecting 1 µL hyphal solution with a laser pulled 10 µL micropipette (Drummond) and aspirator tube (Drummond) into the thorax underneath the front legs. Sham treatments were done in similar fashion using 1 µL medium without hyphae (23).

### Treatments and individual identification

Subcolonies were made of fifteen healthy, ten injected with Graces + FBS media used for *Ophiocordyceps* growth in the laboratory and another ten were injected with *Ophiocordyceps* plus media. These individuals were collected by their colonies by agitating the housing tubes within each colony had and collect the individuals that from the population. In order to follow individuals through time we used a dot system, each individual had a different dot pattern painted on its body. We used an edding^®^ 751 paint marker to label the ants we used for the experiment.

### Behavioral observation set up

We created sub-colonies containing 35 worker ants within a wooden cage with a volume of 14.93 ± 0.53 cm^3^. In order to make 24 hour observations we used a Go Pro camera (Hero 2 with IR lens) and an IR lamp was used for nocturnal observations. The camera was located on top of the colony chamber and removed to change size video card three times a day.

### Trophallaxis

There was only one observer who made the observations of trophallaxis to reduce observer bias. Trophallaxis was classified as starting when labrum was exposed and distended between the two individuals. The event was as over when the mouth parts separated and the individual parted ways. We observed a total of 976.24 hours of video for Colony 2 in order to determine the amount of trophallaxis focal individuals were receiving on days 3-9,12,15 and 18 in trial one. A total of seven Infected individuals and five sham treated and five healthy ants were followed over the course of the daylight session (7.68 ± 0.32 hours per day) on days 3,4,5,6,7,8,9,12,15 and 18 post injection. We analyzed days 3-9 since these are the days we have most infected individuals inside the nest (Figure 1). The chambers in which ants were placed did not restrict individuals to stay within the nest, we were only able to record behaviors for those present within the nest at the time of observation.

### Aggression

There was only one observer who took note of aggressive behavior to reduce observational bias. The videos were observed in fast forward and stopped if any abnormal behavior occurred. Colony 2 has a total of 76.77 hours observed and no aggression has been seen. Further observations will be made in other colonies to see if non-aggression holds.

### Distance data collection

Screen shots were made for every ten minutes of observation during the day period (8.24 ± 0.34 hours per day). Individuals were identified using paint marks. We then used an R program (version 2.15.1; created by Kezia Manlove) that calculated both pair-wise distances and x-y coordinates for the individuals within the chamber. On average there were 24 ± 2 individuals inside the chamber, we recorded point distances on all the individuals visible to us on days 3, 6, 9 and 12 for a total of 4,758 x-y coordinates and 61,000 pair-wise data points.

### Distance data analysis

We focused on days 3, 6 and 9 (8.09 ± 0.43 hours per day) when analyzing the data. We used point process models, which take a set of point locations in window of space. Each of these points can be labeled with a mark to indicate a certain type or class. In this case, the ant locations are the points of interest, the window of observation is the nest, and the mark of each ant is their infection status; either untreated, infected, or sham treated. In point process statistics, the interaction between points can be measured using a summary statistic called the K-function (14, 15). For a given distance *d*, the K-function gives the expected number of additional ants to be found within a radius of *d* of a focal ant.

Functions to compute the K-cross function from healthy to infected ants, and the nearest neighbor to each ant at each time point, were created in R. The K-cross function finds the average number of infected ants within a specified distance of a healthy ant, with the average being over all healthy ants in the chamber at each time point, and over all time points within the specified day. The permutation tests for the nearest-neighbor analysis were carried out by permuting the labels (healthy, infected, or control) of ants in the chamber at each time point and re-computing the nearest neighbor of each ant.

## Acknowledgments

This material is based upon work supported by the National Science Foundation Graduate Research Fellowship under Grant No. DGE1255832. Any opinion, findings, and conclusions or recommendations expressed in this material are those of the authors(s) and do not necessarily reflect the views of the National Science Foundation. We would also like to acknowledge the work done by undergraduate students in the Hughes lab for their data collection.

